# Precision phenotyping reveals novel loci for quantitative resistance to septoria tritici blotch in European winter wheat

**DOI:** 10.1101/502260

**Authors:** Steven Yates, Alexey Mikaberidze, Simon G. Krattinger, Michael Abrouk, Andreas Hund, Kang Yu, Bruno Studer, Simone Fouche, Lukas Meile, Danilo Pereira, Petteri Karisto, Bruce A. McDonald

**Affiliations:** Molecular Plant Breeding, Institute of Agricultural Sciences, ETH Zurich, Zurich, Switzerland; Plant Pathology, Institute of Integrative Biology, ETH Zurich, Zurich, Switzerland; Biological and Environmental Science and Engineering Division, King Abdullah University of Science and Technology (KAUST), Thuwal, Saudi Arabia; Crop Science, Institute of Agricultural Sciences, ETH Zurich, Zurich, Switzerland

**Keywords:** automated image analysis, genome-wide association study (GWAS), plant breeding, precision phenotyping, septoria tritici blotch, *Zymoseptoria tritici*

## Abstract

- Accurate, high-throughput phenotyping for quantitative traits is the limiting factor for progress in plant breeding. We developed automated image analysis to measure quantitative resistance to septoria tritici blotch (STB), a globally important wheat disease, enabling identification of small chromosome intervals containing plausible candidate genes for STB resistance.
- 335 winter wheat cultivars were included in a replicated field experiment that experienced natural epidemic development by a highly diverse but fungicide-resistant pathogen population. More than 5.4 million automatically generated phenotypes were associated with 13,648 SNP markers to perform a GWAS.
- We identified 26 chromosome intervals explaining 1.9–10.6% of the variance associated with four resistance traits. Seventeen of the intervals were less than 5 Mbp in size and encoded only 173 genes, including many genes associated with disease resistance. Five intervals contained four or fewer genes, providing high priority targets for functional validation. Ten chromosome intervals were not previously associated with STB resistance.
- Our experiment illustrates how high-throughput automated phenotyping can accelerate breeding for quantitative disease resistance. The SNP markers associated with these chromosome intervals can be used to recombine different forms of quantitative STB resistance that are likely to be more durable than pyramids of major resistance genes.

## Introduction

Genome-wide association studies (GWAS) provide a powerful approach to identify genetic markers associated with important quantitative traits in crops (e.g. Milner et al. 2018; Yano et al. 2016). The single nucleotide polymorphism (SNP) markers significantly associated with a trait in a GWAS can be directly used in breeding programs for marker-assisted selection or genomic selection, and also as tools to enable map-based cloning of the corresponding genes underlying quantitative traits.

An abundant supply of SNP genetic markers is now available for the most important crops as a result of rapid advances in sequencing technologies. Because phenotyping technologies have not developed as quickly as genotyping technologies, the ability to generate accurate and reproducible phenotypes for quantitative traits is now the primary limitation to progress in breeding for many important traits (Furbank and Tester 2011; Araus and Cairns 2014), including resistance to pests and pathogens (Joalland et al. 2018). Many research teams are working to develop automated and high-throughput phenotyping of important traits under field conditions, with some reports of success (Joalland et al. 2018; Wedeking et al. 2017), but we remain far from the goal of using automated phenotyping to speed progress in plant breeding for useful traits.

Septoria tritici blotch (STB), caused by the fungus *Zymoseptoria tritici*, is currently the most damaging leaf disease on wheat in Europe (Jørgensen et al. 2014) and is a significant disease on wheat around the world. *Z. tritici* has a mixed reproductive system, producing airborne ascospores through sexual reproduction that can be disseminated over distances of several kilometers and asexual conidia that are splash-dispersed over spatial scales of only 1–2 meters over the course of a growing season (McDonald and Mundt, 2016; P. Karisto and A. Mikaberidze, personal communication). *Z. tritici* populations are highly variable within fields as a result of its mixed reproductive system, large effective population sizes and high levels of gene flow among populations (McDonald and Mundt, 2016; Zhan et al. 2003). These properties provide a high evolutionary potential that leads to rapid development of virulence against resistant cultivars (McDonald and Mundt, 2016; Cowger et al. 2000) as well as resistance to fungicides (McDonald and Mundt, 2016; Estep et al. 2015; Estep et al. 2016). STB in Europe is controlled mainly by applying fungicides costing over $1 billion per year (Torriani et al. 2015), but many European *Z. tritici* populations have now evolved sufficiently high levels of resistance that fungicides are losing their efficacy (Karisto et al. 2018; Cools and Fraaije, 2013). The European Union is planning to ban many fungicides during coming years (EU Regulation 1107/2009). These developments have stimulated new efforts to increase STB resistance through plant breeding.

Many studies have identified strain-specific STB resistance genes that could prove useful in breeding programs (summarized in Brown et al. 2015). STB resistance is mainly quantitative, but some examples of major gene resistance were identified (e.g. *Stb6*) that were recently shown to follow the gene-for-gene (GFG) pattern of inheritance (Saintenac et al. 2018; Zhong et al. 2017). Unfortunately, major STB resistance genes like *Stb6* typically failed within 3–4 years of deployment as a result of pathogen evolution (Cowger and Mundt 2000). A different breeding approach that is expected to slow pathogen evolution and be more durable is to make pyramids of quantitative resistance (QR) genes with additive effects (Mundt 2018). This approach requires the identification and deployment of QR that is effective across a broad cross-section of the *Z. tritici* population as opposed to major gene resistance that works against only a small fraction of the strains found in natural field populations.

Identification of QR is difficult for most pathogens for many reasons including: 1) measurement error associated with eyeball assessments of disease; 2) inherent differences in disease measurements conducted by different people; 3) differences in expression of QR in different environments; 4) the occurrence of mixed infections by several pathogens under typical field conditions, with overlapping symptoms that often cannot be teased apart (e.g. STB symptoms look very similar to the symptoms associated with tan spot and stagonospora nodorum leaf blotch). These factors combine to create a low heritability for QR that slows progress in accumulating different sources of QR in breeding programs.

The recent development of automated image analysis for STB enabled rapid acquisition of large datasets including millions of phenotype datapoints that were highly informative under both greenhouse and field conditions, and facilitated the cloning of genes encoding several avirulence effectors, including *AvrStb6* (Zhong et al. 2017) and *Avr3D1* (Meile et al. 2018; Stewart et al. 2018), as well as the *Zmr1* gene affecting melanization of *Z. tritici* colonies and pycnidia (Krishnan et al. 2018; Lendenmann et al. 2014). We took advantage of the high levels of fungicide resistance in Swiss populations of *Z. tritici* by using fungicide treatments to eliminate competing pathogens in a replicated field experiment (Karisto et al. 2018). The fungicide treatments enabled a pure-pathogen readout of quantitative resistance to STB caused by a genetically diverse, natural population of *Z. tritici* in an epidemic that developed under natural field conditions. Here we use this extensive phenotype dataset in a GWAS to identify 26 chromosome intervals associated with quantitative STB resistance in a broad panel of 335 elite European winter wheat cultivars. Many of these intervals explained 6%–10% of the variance for the associated resistance trait. Several of the intervals contained a relatively small number of annotated genes, including genes known to be associated with disease resistance in wheat or other plants. There was a significant enrichment (*P*<0.0001) for genes encoding putative receptor kinases and kinases within the 17 chromosome segments spanning less than 5 Mbp. Other candidate genes for STB resistance encoded NB-LRR proteins, F-box LRR proteins, sugar transporters, an ABC transporter, superoxide dismutase, and a TCP transcription factor, illustrating how automated image analysis can lead to identification of plausible candidate genes for quantitative disease resistance.

## Materials and Methods

335 European winter wheat cultivars chosen from the GABI wheat panel (Kollers et al. 2013) were grown in 1.1 × 1.4 m plots replicated twice as complete blocks at the Field Phenotyping Platform of the ETH research station in Lindau, Switzerland (Kirchgessner et al. 2017). The plots received full agrochemical inputs typically associated with intensive wheat cultivation in Europe, including mineral fertilizers, a stem shortener and several pesticide applications. Among the pesticides, fungicides comprising five different active ingredients with three modes of action were applied at three time points over the growing season. Additional details associated with the field experiment are given in Karisto et al. (2018).

An unusual feature of this experiment is that all STB infection was natural, with the epidemic caused by a highly diverse *Z. tritici* population that immigrated into the experimental plots via windborne ascospores coming from nearby wheat fields that were treated with fungicides. This local *Z. tritici* population carried sufficient resistance to all fungicides applied in the experimental plots to enable an STB epidemic to develop despite the intensive fungicide treatments. But other wheat diseases common in this region, including leaf rust, stripe rust, stagonospora nodorum blotch, powdery mildew and tan spot, were practically absent because the fungicides excluded these pathogens (Karisto et al. 2018). As a result, we were able to obtain a pure-culture read out of quantitative STB resistance across all 335 wheat cultivars without confounding effects from other diseases. The local weather during the 2015–2016 growing season was cooler and wetter than usual, providing a highly conducive environment for development of an STB epidemic. At least six asexual reproduction cycles occurred during the most active period of wheat growth between March and July (Karisto et al. 2018). Other components of STB epidemiology associated with this experiment were already reported (Karisto et al. 2018).

All experimental plots were assessed for STB resistance at two time points, *t*_1_ (20 May 2016, approximately GS 41) and *t*_2_ (4 July 2016, approximately GS 75–85) using automated image analysis of 21,420 scanned leaves infected by *Z. tritici* (Stewart et al. 2016; Karisto et al. 2018). Nearly sixteen infected leaves collected from the same leaf layer in each plot were mounted on A4 paper and scanned at 1200 dpi using flatbed scanners as described earlier (Karisto et al. 2018). The scanned images were analyzed using an ImageJ macro script (Karisto et al. 2018). Automatically generated outputs of the script included percentage of leaf area covered by lesions (PLACL), average pycnidia density within lesions (*ρ*_lesion_), and average pycnidia darkness (measured using the 256-point gray scale). To measure pycnidia sizes, we developed a Python (version: 3.6.7, https://www.python.org/) program based on the determination of contours of constant brightness in the vicinity of each detected pycnidium with the help of the skimage package (version: 0.13, https://scikit-image.org/.) Each of these STB-associated phenotypes were analyzed separately in a GWAS. The grand means for each phenotype were calculated based on an average of 60 scanned leaves for each wheat cultivar, including both time points and both replicates for each plot (i.e. four measurements of each trait associated with STB resistance). The mean values of PLACL and *ρ*_lesion_ were 1/x transformed to better fit a normal distribution, yielding a *P* < 0.01 for the Shapiro-Wilk test after transformation.

The SNP markers used for the GWAS came from the Illumina 90K SNP array (iSELECT, San Diego, USA, Wang et al., 2014). The majority of the markers on this array were not useful for our experiment because they were not polymorphic in the GABI panel. The remaining markers were positioned on the IWGSC wheat genome (IWGSC, 2018) using a BLASTN search with E-value < 10^−30^. The position with the lowest E-value was assigned as the marker position. In the case of ties where it was not possible to unequivocally assign a marker to one of the homeologous chromosomes, the markers were omitted. Additional filtering criteria to choose SNPs for the GWAS were: a call rate of > 95 % per marker, > 5 % minor allele frequency and identity by state (IBS) < 0.975, using the GenABEL software in the R statistical environment (Aulchenko *et al*. 2007). After filtering, a total of 13,648 high quality SNP markers were used for the GWAS. Haplotypes were identified using a sliding window of three consecutive SNPs with PLINK (Purcell *et al*. 2007) and tested using linear regression models. GWAS Manhattan plots were constructed using R (version 3.5.1, R core team, 2018) with ggplot2 (version: 3.1.0. Wickham, 2016). Bonferroni thresholds were calculated using *P*/N (0.05/13,648) yielding a LOD (-log_10_(*P*)) score of 5.44. The fraction of the phenotypic variance associated with the 26 chromosome intervals at or exceeding the Bonferonni threshold was calculated using linear regression models in R (*lm* function). The adjusted *R^2^* provided a measure of the proportion of the variance explained.

The coordinates of the 26 intervals exceeding the Bonferonni threshold were plotted onto the Chinese Spring reference genome as described earlier and used to compare the positions of the SNPs affecting STB resistance identified in this analysis with the positions of STB resistance traits identified in earlier studies (Brown et al. 2015). The sequence data of the markers associated with STB resistance in earlier studies were retrieved from GrainGenes (https://wheat.pw.usda.gov/GG3/) and then searched using BLASTN against the IWGSC 2018 assembly using Unité de Recherche Génomique Info (URGI, https://wheat-urgi.versailles.inra.fr/) for the corresponding chromosome. The position of the best hit was used as the genome position.

Candidate gene identification was based on the gene annotation of the IWGSC v1.0 reference sequence of the wheat landrace Chinese Spring (IWGSC 2018). All high confidence genes in chromosome segments shorter than 5 Mb were identified.

## Results

The three fungicide treatments eliminated all competing fungal pathogens, enabling a mono disease readout of the relative degree of quantitative resistance to STB under the field conditions typically used for intensive wheat production in Europe. All STB infection was natural, with an epidemic resulting from at least six cycles of infection by a diverse *Z. tritici* population that included a high degree of gene and genotype diversity, with infections caused by millions of different *Z. tritici* strains despite the fungicide applications. As an indicator of the pathogen genetic diversity in these plots, genome sequences of 161 *Z. tritici* isolates obtained from 21 of the plots revealed 147 unique genome sequences, with all but two of the identified clones found within the same 2 m^2^ plot (Daniel Croll, personal communication). This high level of pathogen diversity was consistent with earlier findings from other naturally infected wheat fields around the world and was expected given that *Z. tritici* populations experience high levels of recombination (Chen and McDonald 1996; Zhan et al. 2003) that enable different fungicide resistance mutations to segregate and re-assort into many different genetic backgrounds in natural populations.

The quantitative measures of STB severity generated by automated image analysis followed the continuous distribution typically associated with quantitative traits (Karisto et al 2018; Stewart et al. 2016). Earlier analyses of relationships among these traits (Karisto et al 2018) showed that resistance that minimizes host damage (PLACL) was largely independent of resistance that minimizes pathogen reproduction (*ρ*_lesion_). Hence the GWAS was conducted independently for each trait. In addition to the traits PLACL and *ρ*_lesion_, we measured the average size of pycnidia formed within lesions, which reflects the average size and number of spores contained in each fruiting body (Stewart et al. 2018), (i.e. pycnidia size is an independent indicator of pathogen reproduction), and the average gray value of pycnidia, which reflects the average amount of melanin accumulated in each fruiting body (Stewart et al. 2018; Krishnan et al. 2018). Our earlier work indicated that pycnidia melanization is on average greater on wheat cultivars with more resistance to STB (Stewart et al. 2016; Stewart et al. 2018). Altogether, the phenotype measurements used for the GWAS included 21,420 measures each of PLACL and *ρ*_lesion_ and 2.7 million measures each of pycnidia size and pycnidia melanization, yielding a total of >5.44 million automatically measured phenotypes that were not prone to human scoring error.

Manhattan plots for PLACL, *ρ*_lesion_, pycnidia size, and pycnidia gray value revealed the SNPs with the highest associations for each STB resistance trait (Figure 1). A total of 109 SNPs were at or above the Bonferroni threshold across all traits based on the GWAS. Marker-trait associations were calculated using sliding windows including three consecutive SNPs. Among these, 52 haplotypes were at or above the Bonferroni threshold. Further evaluation of the 52 haplotypes revealed overlaps that were combined to produce a non-redundant set of 26 chromosome segments that explained from 1.9% to 10.6% of the overall variance associated with each resistance trait (Table 1).

**Figure 1.**
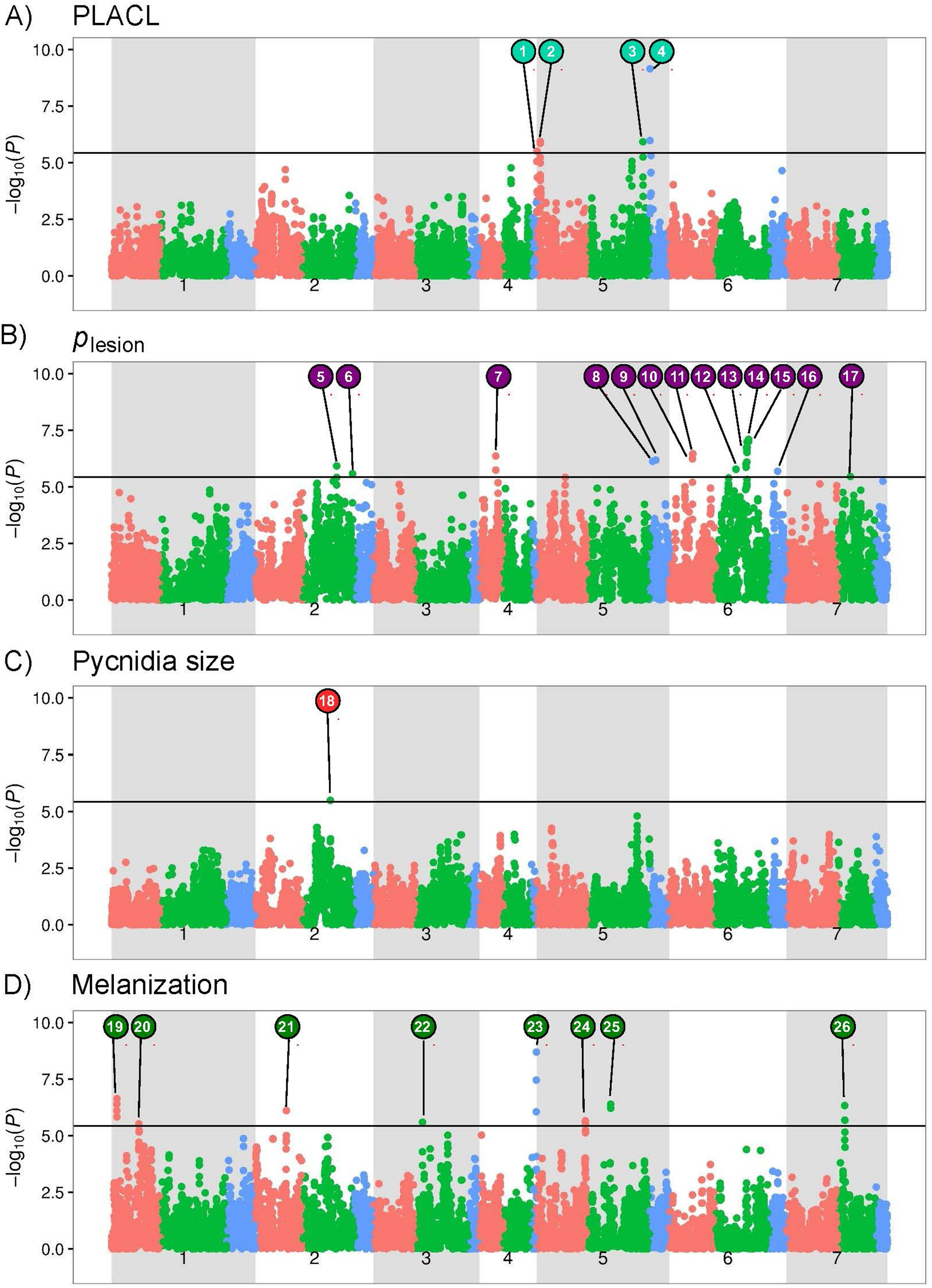
Manhattan plots showing significant SNP markers associated with each trait. The horizontal line indicates the Bonferroni-adjusted significance threshold. The A, B and D genomes of wheat are shown in red, green and blue, respectively. SNPs associated with the interval IDs shown in Table 1 are indicated in colored circles. A. Percentage of leaf area covered by lesions (PLACL) had four significant associations distributed across chromosomes 5A, 5B and 5D. B. Density of pycnidia within lesions (*ρ*_lesion_) had 12 significant associations distributed across chromosomes 2B, 4A, 5A, 5D, 6A, 6B, 6D and 7B. C. Pycnidia size had a single significant association located on chromosome 2B. D. Pycnidia melanization had 8 significant associations distributed across chromosomes 1A, 2A, 3B, 4D, 5A, 5B, and 7B.

For the PLACL trait that reflects the ability of a wheat cultivar to limit the degree of necrosis caused by an STB infection, 14 SNPs identified 4 different genomic positions distributed across chromosomes 5A, 5B and 5D with LOD scores exceeding 5.5. Interval 4 on 5D had a LOD score of 9.2 and explained 10.3% of the total variance associated with PLACL (Table 1). For the *ρ*_lesion_ trait that reflects the ability of a wheat cultivar to restrict *Z. tritici* reproduction, 51 SNPs identified 13 genomic positions located on chromosomes 2B, 4A, 5D, 6A, 6B, 6D and 7B, with LOD scores ranging from 5.5 to 7.1. Interval 15 on 6B had the highest LOD score and explained 9.3% of the total variance associated with *ρ*_lesion_. For the pycnidia size trait, three SNPs located on 2B defined a chromosome interval that surpassed the Bonferroni threshold. Interval 18 explained 5.9% of the total variance associated with pycnidia size. For the pycnidia melanization trait, 36 SNPs defined 8 genomic positions located on chromosomes 1A, 2A, 3B, 4D, 5A, 5B, and 7B. Interval 23 on 4D showed the highest LOD score of 8.7 and explained 10.6% of the total variance associated with pycnidia melanization.

The positions of the 26 chromosome segments identified in this experiment were compared to the positions of mapped STB genes reported in earlier publications (summarized in Fig. 1 of Brown et al. 2015). Figure 2 shows that 16 of the 26 chromosome segments identified in our analyses overlapped with or were very close to genomic regions identified in earlier publications, while 10 of the chromosome segments were in chromosomal regions that were not previously associated with STB resistance. Among those, two were associated with PLACL (3, 4), five with *ρ*_lesion_ (7, 8, 9, 16, 17), and three with pycnidia melanization (20, 23, 26) (Table 1).

**Figure 2.**
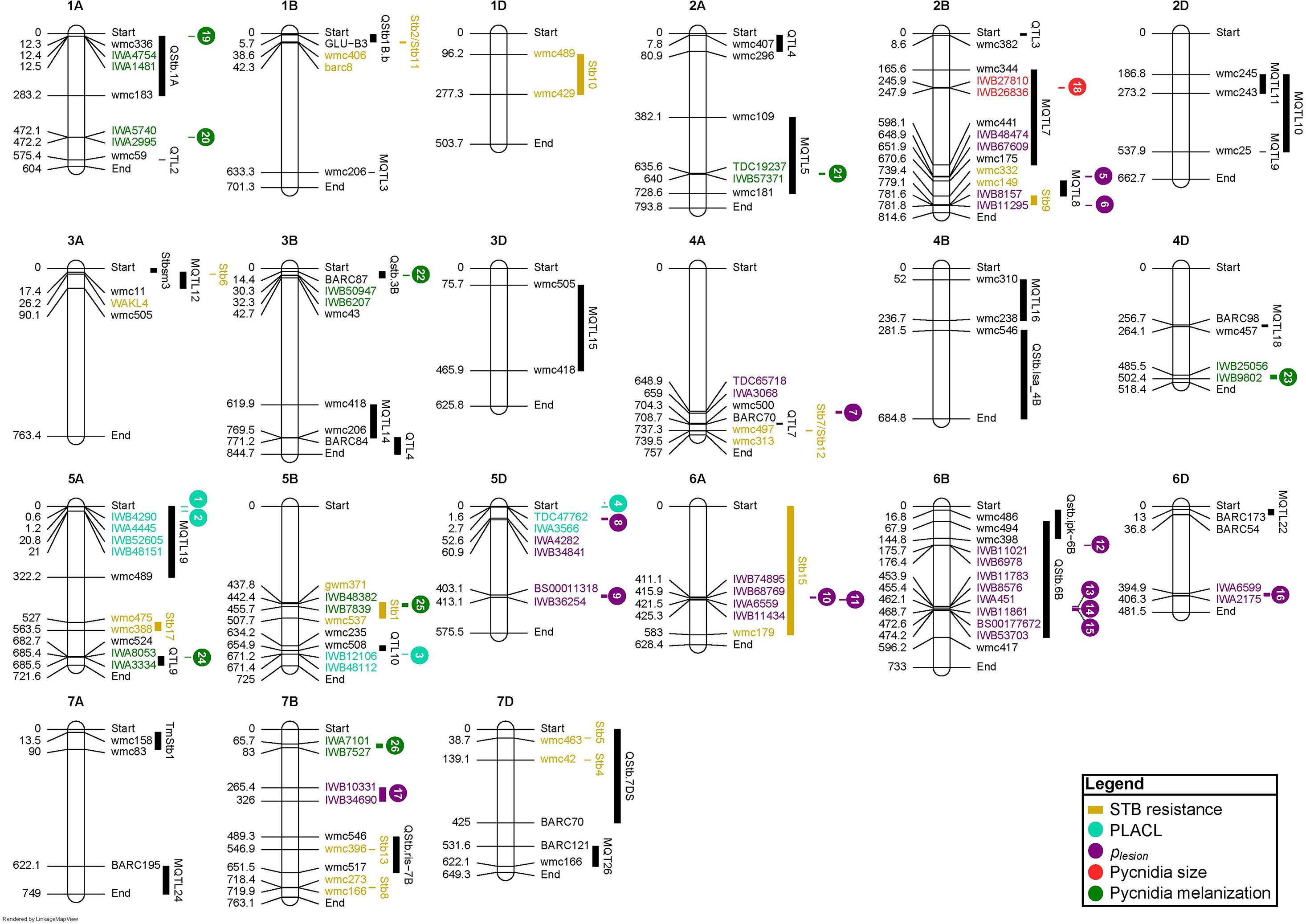
Positions on the Chinese Spring reference genome (IWGSC 2018) of 26 significant GWAS marker-trait associations across four resistance traits compared to positions of previously mapped STB resistance genes (Brown et al. 2015). The 26 associations are shown as numbered circles and a bar (95% confidence interval) in cyan for PLACL, purple for *ρ*_lesion_, red for pycnidia size and green for melanization. Confidence intervals of previously mapped STB resistance loci are shown in dark blue bars (STB genes) and black bars (QTLs). SNP markers are presented as locus names from GrainGenes (https://wheat.pw.usda.gov/GG3/) for brevity. Markers with the prefix Tdurum_contig were abbreviated to TDC. Only SNP markers with significant associations with STB genes, QTLs or the four phenotypes included in the GWAS are shown. For each association confidence interval, the first and the last SNP and their positions are shown. Names are colored according to the type of association.

Seventeen of the 26 chromosome segments were smaller than 5 Mb. For these, we identified putative candidate genes responsible for STB resistance based on the wheat reference genome sequence of Chinese Spring (IWGSC 2018). In total, the 17 intervals spanned 24.2 Mb and contained 173 high confidence genes (Supplementary Table 1). There was a significant enrichment (*P*<0.0001) for genes encoding putative receptor kinases and kinases within these 17 chromosome segments. Receptor kinase genes were recently shown to play major roles in disease resistance in cereals (Saintenac et al. 2018; Keller and Krattinger 2018; Ma et al. 2018), including the *Stb6* gene encoding resistance to STB (Saintenac et al. 2018). Five of the chromosome segments contained four or fewer genes, with three of these segments (19, 20, 24) associated with pycnidia gray value and two segments (2, 3) associated with PLACL. The smallest chromosome segment (20) encompassed 28 kb on chromosome 1A and contained a single gene in Chinese Spring (TraesCS1A01G277000) encoding a putative solute carrier family 35 member. The 99.7 kb segment 24 on the long arm of chromosome 5A also had a single gene (TraesCS5A01G524800) encoding a putative 4-hydroxy-tetrahydrodipicolinate reductase, a protein involved in lysine biosynthesis. Intervals 2 (chromosome 5A) and 19 (chromosome 1A) had three candidate genes each, of which a putative kinase gene and a putative nucleotide binding site – leucine-rich repeat (NLR) represent the most obvious candidates as these categories of genes are known to affect disease resistance (Dodds and Rathjen, 2010). Interval 3 contained four candidate genes, all of which were associated with F-box proteins, a class of proteins often associated with plant defense responses (van den Burg et al. 2008). Other candidate genes known to be involved in disease resistance include sugar transporters (associated with PLACL in intervals 1 and 4), superoxide dismutase (associated with pycnidia size in interval 18), an ABC transporter (associated with PLACL in interval 1), and a TCP transcription factor (associated with *ρ*_lesion_ in interval 15).

## Discussion

In a year that was highly conducive to development of an STB epidemic, we combined a novel automated image analysis tool that could differentiate independent components of STB severity with the high level of fungicide resistance existing in a local Swiss population of *Z. tritici* to make a quantitative comparison of STB resistance across a broad cross section of elite European winter wheat cultivars. GWAS analyses that coupled these quantitative measures of STB resistance with 13,648 SNP markers enabled identification of 109 SNPs on 13 chromosomes that defined 26 chromosome segments highly associated with STB resistance. Because all STB infection in this experiment was natural, including millions of different pathogen genotypes originating from a recombining population, and the growing season was highly conducive to development of an STB epidemic, we believe that the SNP markers defining the chromosome intervals associated with the highest levels of STB resistance could be especially useful in European breeding programs aiming to increase overall levels of STB resistance to the *Z. tritici* populations found in Europe.

The 26 chromosome intervals associated with STB resistance ranged from 28 kbp to 60 Mbp in size and were distributed across 13 chromosomes, with individual intervals explaining 1.9% to 10.6% of the phenotypic variance for each trait. Some of the intervals were clustered in the same chromosomal region (e.g. intervals 1 and 2 associated with PLACL on chromosome 5A; intervals 13, 14 and 15 associated with pycnidia gray value on chromosome 6B), but most of the intervals were genetically distant from each other. Sixteen of the intervals were embedded within or located very close to chromosomal regions previously associated with STB resistance, but 10 intervals were in genomic regions that had not been associated with STB resistance. Particularly notable novel regions were the intervals 4, 8 and 9 located on 5D, a chromosome which had not previously been associated with STB resistance (Brown et al. 2015), though Kollers et al (2013) found some weak associations with STB resistance on this chromosome. There was no overlap between chromosomal segments associated with host damage (PLACL) and pathogen reproduction (pycnidia density or pycnidia size), indicating that these resistance traits were under independent genetic control as hypothesized earlier (Karisto et al. 2018). Intervals 4, 14, 15, and 23 had LOD scores at or exceeding 7. These are candidate regions for genes encoding broadly based field resistance to STB that may be especially useful against the genetically diverse *Z. tritici* populations in Europe.

The chromosome segments identified in our GWAS are much smaller than the intervals defined in earlier work as shown in Figure 2. For example, STB15 was previously mapped to a region that includes most of chromosome 6A (∼590 Mbp) while we identified two separate chromosome regions (10 and 11) within the STB15 region that encompass only ∼8.5 Mbp. Similarly, STB1 was mapped to a region that covered ∼69.9 Mbp on chromosome 5B while interval 25 covers only ∼13 Mbp within this region. The smaller intervals detected in our GWAS reflects the much higher marker density used in our experiment coupled with more accurate knowledge of marker positions coming from the new wheat genome assembly. Other contributors to the small intervals were the more accurate quantitative phenotypes yielding relatively large effect sizes and the haplotype-based GWAS approach that increased the statistical power compared to standard GWAS pipelines.

Seventeen of the 26 chromosome segments identified in the GWAS were less than 5 Mbp in size and contained between 1–28 candidate genes annotated in the Chinese Spring reference genome. The 173 genes located in these intervals were significantly enriched for receptor kinases and kinases, including clusters of 6 and 10 kinases found in intervals 18 and 22 respectively. We consider this enrichment to be notable because *Stb6*, the only cloned STB resistance gene, is a receptor kinase (Saintenac et al. 2018). Also notable was our finding that genes encoding receptor kinases are strongly upregulated during infection by all tested strains of *Z. tritici* (Ma et al. 2018). Hence we hypothesize that some of the receptor kinase genes found in these intervals may be responsible for the STB resistance we observed. The interval 4 associated with PLACL explained 10.3% of the overall variance and provided the first report of STB resistance on chromosome 5D. This interval contained 12 genes, including three encoding proteins already shown to affect disease resistance, including an NLR, a S/T protein kinase and a sugar transporter (Dodds and Rathjen 2010; Moore et al. 2015). We hypothesize that one or more of these genes are responsible for the STB resistance in this chromosome segment. Other interesting candidate genes found in the 17 intervals encode an ABC transporter, a TCP transcription factor and superoxide dismutase. The *Lr34* gene encoding quantitative resistance to leaf rust and other diseases in wheat was shown to be an ABC transporter (Krattinger et al. 2009). Superoxide dismutases are involved in synthesis of hydrogen peroxide, which was already shown to be involved in wheat’s defense response against STB (Shetty et al. 2007). TCP transcription factors were shown recently to be important components of the signaling pathway involved in systemic acquired resistance (Li et al. 2018). Segment 24, which explained 8% of the variance in pycnidia melanization and lies within the QTL9 region identified in earlier mapping studies, contained a single gene encoding a protein involved in lysine biosynthesis. Recent work on the wheat pathogen *Cochliobolus sativus* showed that lysine was essential for melanin biosynthesis (Leng and Zhong 2012) and lysine was recently shown to be essential for virulence in *Z. tritici* (Derbyshire et al. 2018). We conclude from this analysis that many of the genes found in the intervals identified in the GWAS are plausible candidates to explain the observed phenotypes associated with STB resistance, but functional validation studies will be needed to confirm whether any of these genes actually play a role in resistance.

Earlier field trials also used association mapping to identify genetic markers associated with STB resistance (Kollers et al. 2013; Miedaner et al. 2013; Muqaddasi et al. 2019). In all of these trials, the experimental plots were inoculated with a small number of *Z. tritici* isolates that were sprayed when all wheat genotypes had fully extended flag leaves (i.e. GS >41) a few weeks before scoring for STB resistance. As a result, the associations identified in those experiments are likely to be strain-specific and represent the outcome of a single cycle of infection based on a high dose of artificially applied blastospore inoculum. Similarly, most experiments that identified STB genes with major effects were based on greenhouse inoculations of seedlings by a single pathogen strain and used disease scores made at a single point in time, leading to identification of genes that encode seedling resistance to the strain used in the experiment. It is now clear that natural field infections of STB are caused by many millions of *Z. tritici* strains, with a different strain occurring on each infected leaf, on average, and with most leaves infected by more than one strain (Linde et al. 2002). The significant STB resistance associations identified in our experiment were based on a natural epidemic that included at least six cycles of pycnidiospore infection by a highly diverse population of the pathogen and included two time points during epidemic development. We believe that the STB resistance identified in our experiment is more likely to be broadly applicable under natural field conditions and hence more useful in breeding programs aiming for stable STB resistance.

An important and novel aspect of our experiment was the use of an automated image analysis pipeline for phenotyping that eliminated human scoring bias while generating millions of accurate phenotype data points. As is the case for many plant diseases (Saari and Prescott 1975), the traditional eyeball assessment of STB typically generates a single number on a 0–9 scale (Eyal et al. 1987) that tries to integrate the totality of disease in a particular plot, often relative to other plots in the same field or trial. Eyeball assessments are fast, often requiring less than one minute per plot to produce a measurement, but are prone to variation caused by fatigue, changes in lighting over the course of a day, and differences in opinion among different scorers. The automated image analyses allowed us to simultaneously assess four quantitative phenotypes that could not be accurately measured by eye. A traditional eyeball assessment would have generated a total of 4 STB measurements per cultivar to use in the GWAS. Our automated analyses generated an average of over 16,000 STB measurements per cultivar. These detailed phenotype data enabled us to separate different components of STB resistance, in particular allowing us to separate STB resistance that affects host damage (PLACL) from STB resistance that affects pathogen reproduction (*ρ*_lesion_ and pycnidia size). We believe that resistance affecting pathogen reproduction is likely to be more effective in the long run for several reasons, including: 1) Our earlier analyses (Karisto et al. 2018) showed that measures of pathogen reproduction (*ρ*_lesion_) early in the growing season were the best predictors of host damage (PLACL) late in the growing season, showing that resistance that reduces pathogen reproduction is likely to minimize yield losses caused by STB; 2) A decrease in pathogen reproduction diminishes the amount of inoculum available to cause new cycles of infection, which will lower the transmission rate (i.e. decrease the basic reproductive number, R_0_) during each infection cycle and result in less overall infection by the end of the epidemic; 3) A decrease in pathogen inoculum will lead to a decrease in the pathogen population size, which will decrease the overall genetic diversity and provide fewer opportunities for favorable mutations (e.g. for fungicide resistance or gain of virulence) to emerge (Stam and McDonald, 2018). This should lower the overall evolutionary potential of the pathogen population (McDonald and Linde, 2002).

Recombining the SNP markers associated with the STB resistance intervals identified in this experiment may accelerate breeding efforts aiming to increase quantitative resistance to STB in European wheat. We showed that resistance affecting leaf damage (PLACL) is genetically distinct from resistance affecting pathogen reproduction (*ρ*_lesion_). We consider it likely that these different resistance phenotypes reflect different underlying mechanisms of STB resistance. We hypothesize that PLACL reflects the additive actions of toxin sensitivity genes that interact with host-specific toxins produced by the pathogen, as shown for *Parastagonospora nodorum* on wheat (e.g. Friesen et al., 2008; Oliver et al., 2012), while pycnidia density reflects the additive actions of quantitative resistance genes that recognize pathogen effectors (e.g. Meile et al. 2018). Under this scenario, breeders should aim to recombine these two forms of resistance into the same genetic background, bringing together different forms of resistance that may be more durable when deployed together than when either mechanism is deployed in isolation. We anticipate that functional analyses of the most compelling candidate genes identified in this experiment will enable us to identify new genes underlying the different STB resistance traits. Our experiment illustrates how high-throughput automated phenotyping can accelerate breeding for quantitative disease resistance.

## Acknowledgements

STB research in BAM’s lab was supported by the Swiss National Science Foundation (grants 155955, 134755, 104145 and 56874) and the ETH Zurich Research Commission (grants 12–03, 15–02). AM and PK were supported by the Swiss National Science Foundation through Ambizione grant PZ00P3_161453. H. Zellweger managed the wheat trial. Marion Roeder from IPK Gatersleben provided seeds and marker information for the GABI wheat panel.

## Author contributions

B.A.M. conceived and managed the project. B.A.M., A.M., and B.S. provided funding for the project. A.H., S.F., L.M., D.P., and P.K. performed the experiments. S.Y., A.M., P.K., S.G.K., M.A., and K.Y. analysed the data. B.A.M., S.Y., and S.G.K. wrote the manuscript.

## Supplementary information

**Table S1.**
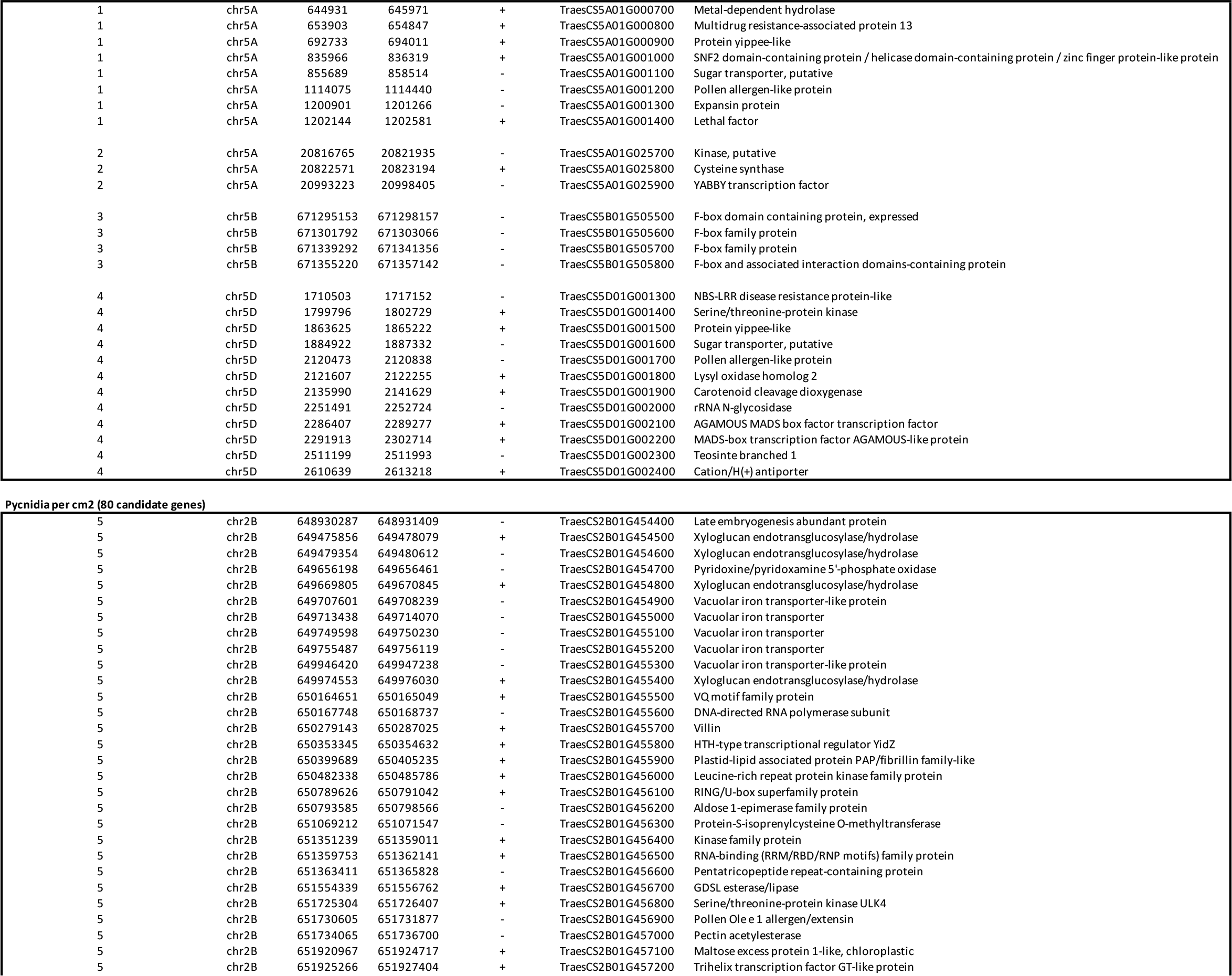
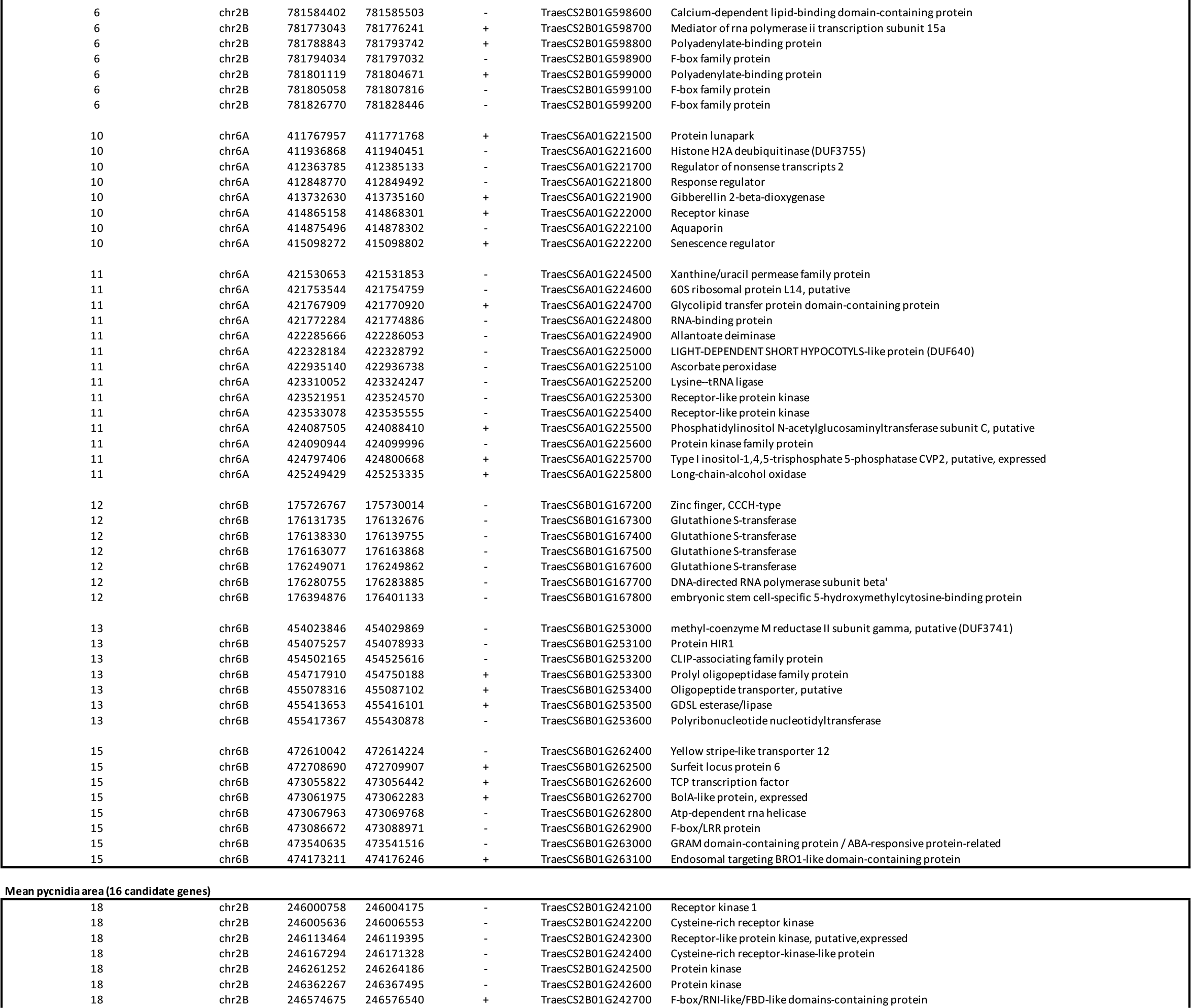
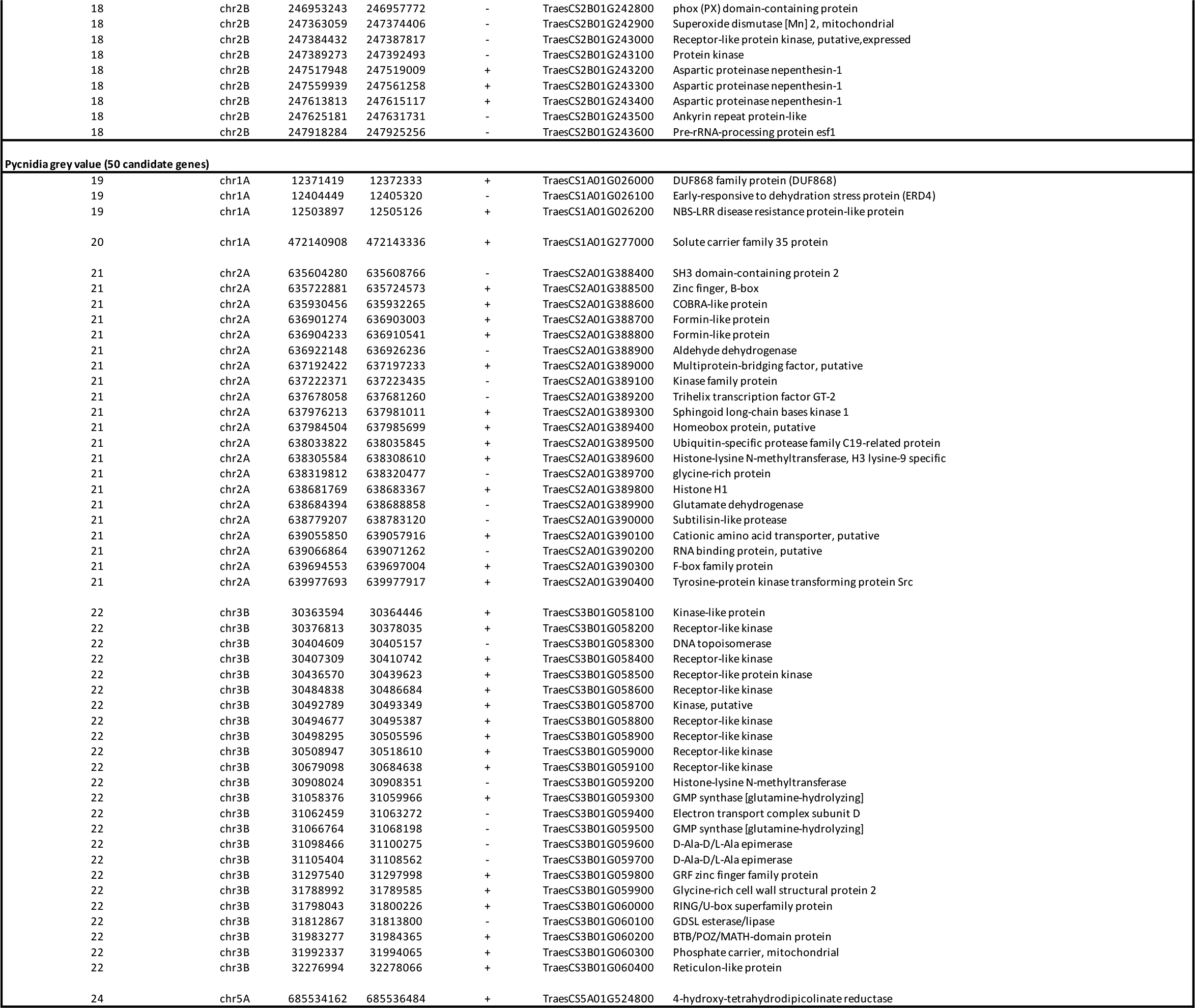
Candidate genes in the chromosome intervals defined by the GWAS.

